## Supporting Information for "Recognition and Cleavage of Human tRNA Methyltransferase TRMT1 by the SARS-CoV-2 Main Protease"

* Jeffrey S. Mugridge

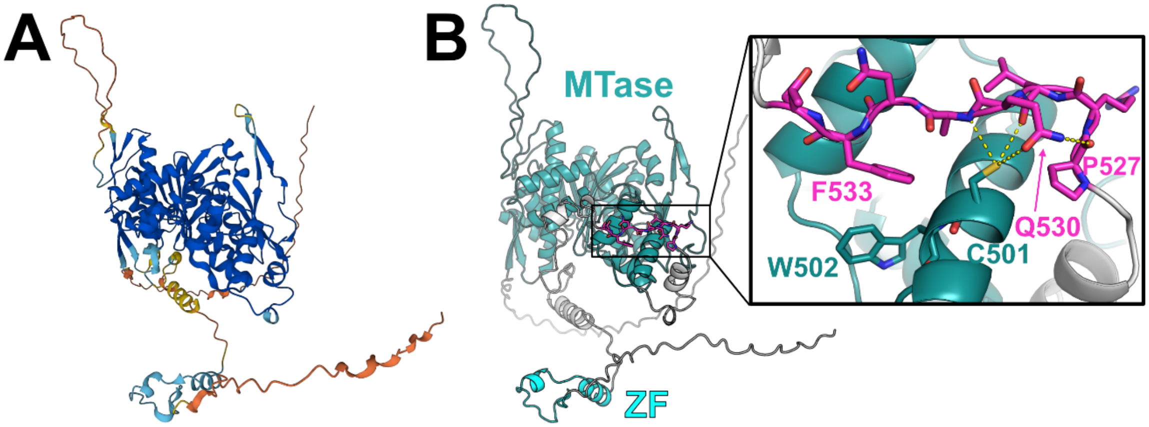

**Figure S1.** AlphaFold-predicted[31,32] structure of human TRMT1. **A**) TRMT1 structural model colored by AlphaFold prediction confidence (dark blue = very high confidence, cyan = confident, yellow = low confidence, orange = very low confidence). **B**) TRMT1 structural model colored by domain (SAM-dependent methyltransferase, MTase = teal; zinc finger, ZF = cyan; linker and unstructured regions = gray) with the TRMT1(527-534) cleavage sequence highlighted in magenta. Inset shows closeup of the surface-exposed TRMT1(527-534) cleavage sequence (magenta) and the AlphaFold-predicted contacts between the residues in the cleavage sequence and the surface of the MTase domain.

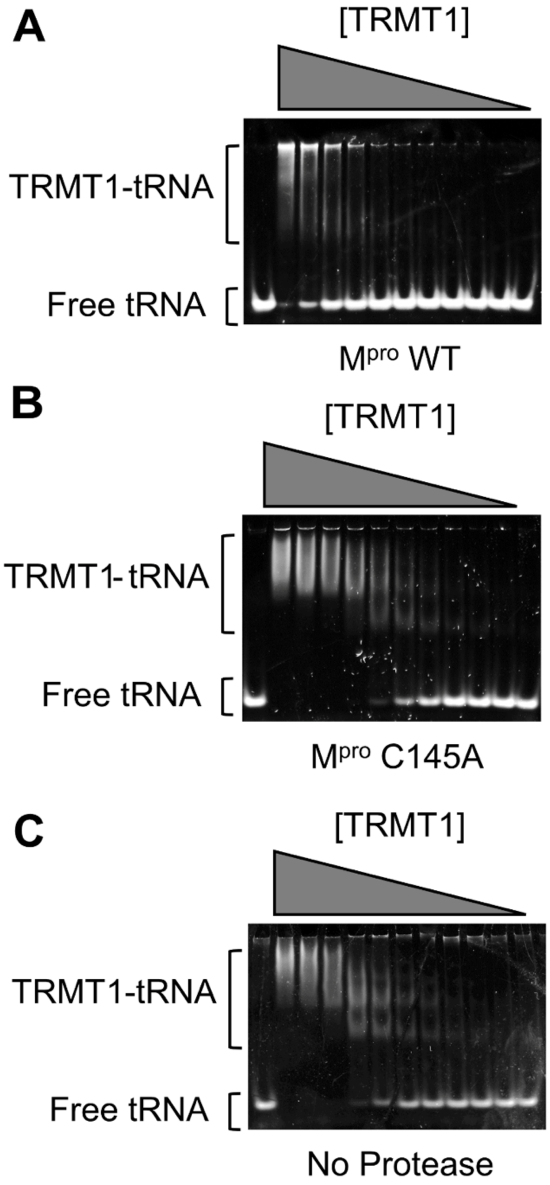

**Figure S2.** TRMT1 binding to tRNA$^{phe}$ substrate was measured using an Electrophoretic Mobility Shift Assay (EMSA). Representative EMSA gels are shown for tRNA binding experiments where TRMT1 was pre-incubated overnight with M$^{pro}$ WT (**A**), M$^{pro}$ C145A (**B**), or no protease (**C**); after overnight treatment with protease (or no protease), binding was measured by EMSA to 300 nM tRNA$^{phe}$, with a TRMT1 dilution series from 7.8 → 0.1 μM. For an unbound tRNA migration reference, the first lane of each gel contained tRNA$^{phe}$ without TRMT1. The cleaved TRMT1 generated by M$^{pro}$ WT (**A**) had diminished affinity for tRNA, whereas TRMT1 incubated with M$^{pro}$ C145A (**B**) had comparable tRNA binding affinity to TRMT1 incubated with no protease (**C**). $K_D$s for the TRMT1-tRNA interactions under each of these conditions are listed in **Table S2** and were determined by quantifying fraction bound from the free tRNA and TRMT1-tRNA bands shown above, plotting this against [TRMT1], and fitting these plots (shown in Figure 2D) to a standard single-site ligand binding equation. We note that the cleaved TRMT1-tRNA complexes in **A** migrate at significantly higher apparent molecular weights, and are retained in the wells, as compared to uncleaved TRMT1-tRNA complexes. Based on our complementary kinetic experiments shown in Figure 2C, although cleaved TRMT1 apparently retains some binding affinity for tRNA, these are non-functional complexes or oligomers.

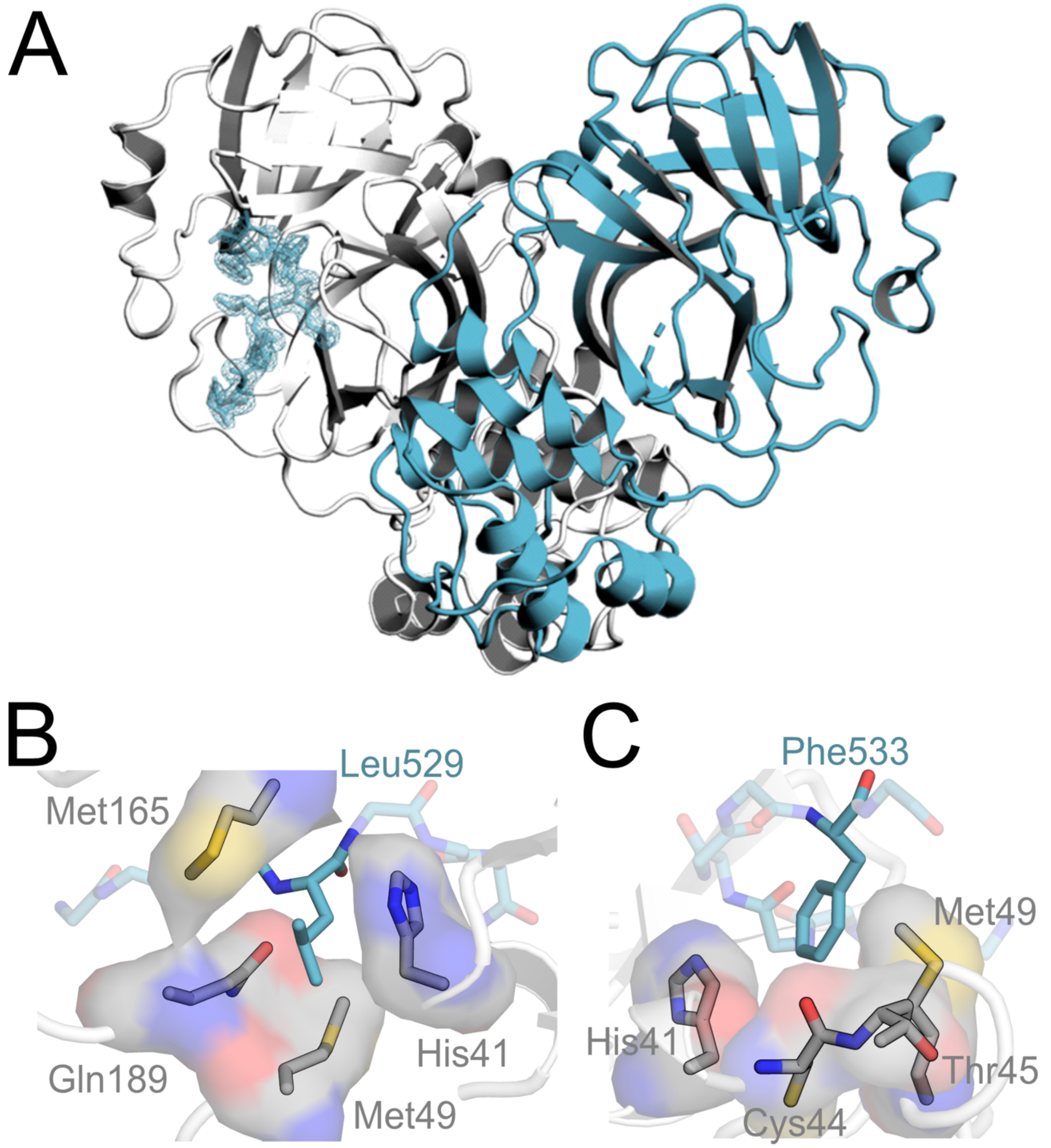

**Figure S3. A)** Structure of human TRMT1(526-536) peptide in complex with the SARS-CoV-2 M$^{pro}$ dimer. Only one protomer of the M$^{pro}$ dimer has TRMT1 bound in the active site. $F_o$-$F_c$ omit electron density map of TRMT1 peptide bound to M$^{pro}$ contoured at 1σ. **B)** TRMT1(526-536) P2 residue Leu529 packs against M$^{pro}$ residues His41, Met49, Gln189, and Met165 in the M$^{pro}$ S2 substrate binding pocket. **C)** TRMT1(526-536) P3′ residue Phe533 sits in the S3′ pocket, composed of M$^{pro}$ residues His41, Cys44, Thr45 (backbone), and Met49.

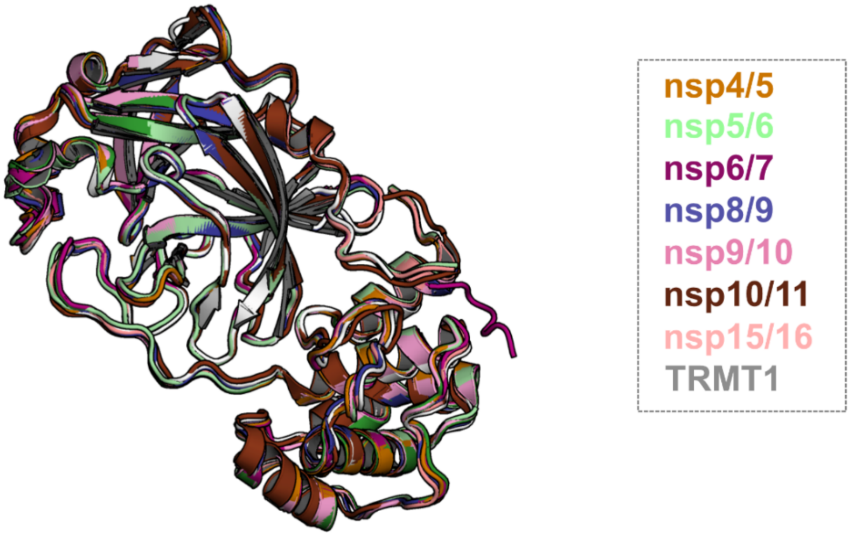

| pdb | $M^{pro}$ Mutation | Peptide Bound | Alignment RMSD |
|---|---|---|---|
| 7T8M | C145A | nsp5/6 | 0.59 |
| 7DVX | H41A | nsp6/7 | 1.52 |
| 7T9Y | C145A | nsp8/9 | 0.75 |
| 7TA4 | C145A | nsp9/10 | 0.70 |
| 7TA7 | C145A | nsp10/11 | 1.07 |
| 7TC4 | C145A | nsp15/16 | 0.53 |
| 8D35 | C145A | TRMT1 | 0.83 |
| **7MGS** | **C145A** | **nsp4/5** | **0.00** |

**Figure S4.** An alignment of $M^{pro}$ structures for each $M^{pro}$-peptide complex used in the analysis shown in Figure 3B and C (**top**). Calculated all-atom RMSDs are derived from each structure alignment to $M^{pro}$ Cys145Ala bound to nsp4/5 (7MGS) (**bottom**). The overall structure of the $M^{pro}$ backbone is highly similar regardless of the bound peptide substrate.

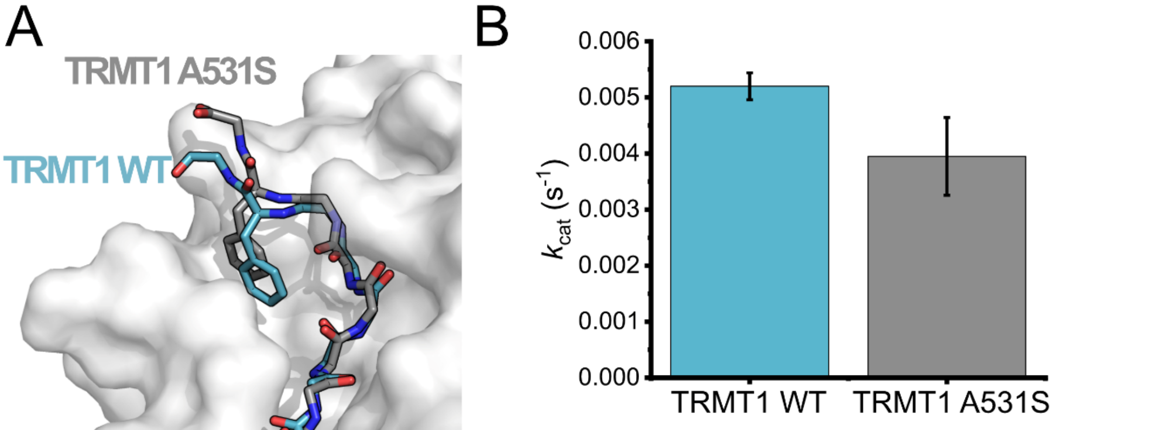

| | P4 | P3 | P2 | P1 | P1′ | P2′ | P3′ | P4′ |
|---|---|---|---|---|---|---|---|---|
| **TRMT1 WT** | P | R | L | Q | A | N | F | T |
| **TRMT1 A531S** | P | R | L | Q | S | N | F | T |

**Figure S5.** Analysis of a point mutation to P1′ Ala residue of TRMT1(526-536). TRMT1(A531S) contains a Ser at position P1′, which conforms to the $M^{pro}$ cleavage consensus (Figure 1B) but is predicted to disfavor the P3′-in binding conformation observed for TRMT1 in the $M^{pro}$ active site. **A)** Molecular dynamics simulated structure of $M^{pro}$ in complex with TRMT1 A531S (gray peptide) predicts this peptide remains in the P3′-in conformation, as observed with TRMT1 WT (blue peptide). **B)** No major difference is measured for cleavage of the WT vs A531S TRMT1 peptide by $M^{pro}$ kinetic data comparing $k_{cat}$ for cleavage of the TRMT1(526-536) peptide with either the WT TRMT1 sequence or an Ala531 to Ser mutation.

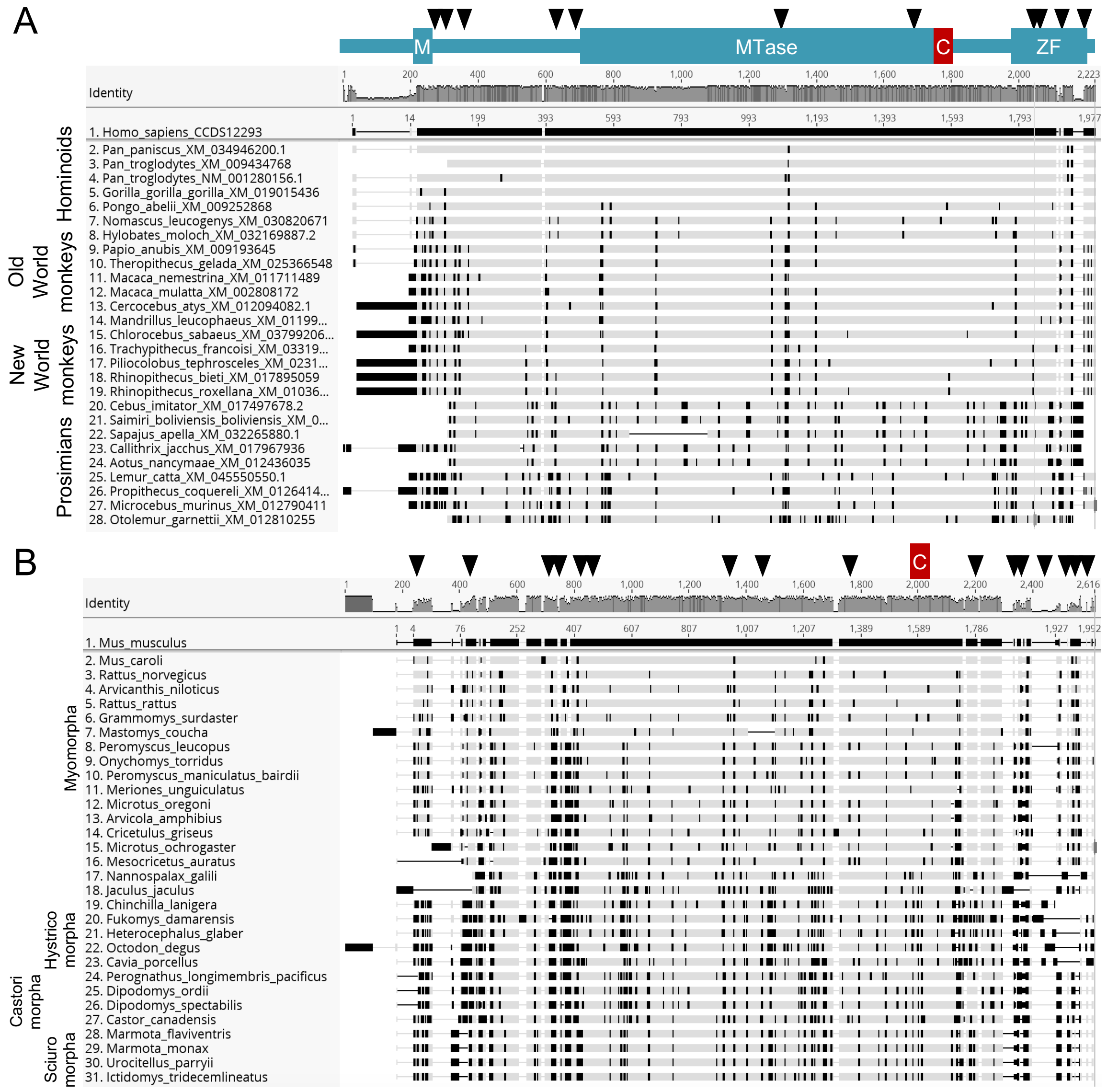

**Figure S6.** Evolution of mammalian TRMT1. **A)** The M^pro binding/cleavage site in TRMT1 is highly conserved in primates, while there has been rapid evolution at N- and C-termini. Codon alignment of the primate TRMT1 sequences with the human as a reference. The numbering is according to the nucleotide position in the alignment (top) and in the reference sequence (human). Non-synonymous differences to the reference are highlighted in black, from Geneious R9. As indicated in Table S3, the sites under positive selection identified by MEME or FUBAR are shown above the alignment with black triangles. Major domains and the cleavage sites are also represented: M for mitochondrial signal, MTase for methyltransferase, C for cleavage by M^pro, ZF for Zinc finger. **B)** Same as panel A for Rodents, with mouse as the reference.

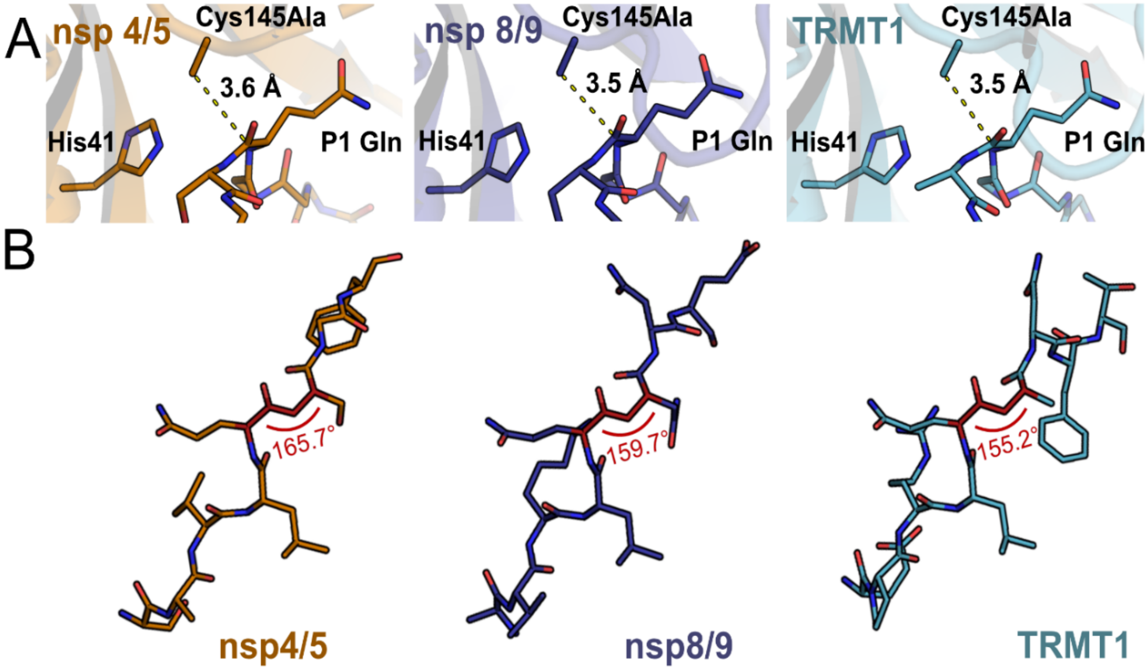

**Figure S7.** Substrate positioning at the M^pro catalytic site does not readily explain observed differences in cleavage kinetics. **A)** One possible explanation for faster cleavage kinetics of nsp4/5 relative to nsp8/9 or TRMT1 (data in Figure 4) could be better positioning of the scissile peptide bond and electrophilic P1 amide carbonyl closer to the nucleophilic M^pro Cys145 residue. However, the measured C145A – P1(CO) distances are nearly identical for nsp4/5-, nsp8/9-, and TRMT1-bound M^pro crystal structures, suggesting this is not the case. **B)** Another possible explanation for faster cleavage kinetics of nsp4/5 are deviations in the dihedral angle of the scissile amide (P1(CA)-P1(C)-P1′(N)-P1′(CA)) bond away from 180°, which could indicate ground state destabilization that would result in accelerated peptide bond cleavage. However, the most rapidly cleaved substrate, nsp4/5, has a scissile amide bond dihedral angle closest to 180°, indicating that amide bond planarity of the bound substrate does not play an important role in determining peptide cleavage rates.

| | Crystal structure of SARS-CoV-2 main protease ($M^{pro}$) in complex with peptide from human tRNA methyltransferase TRMT1 (PDB 9DW6) |
|---|---|
| **Data collection** | |
| Space group | P 21 21 21 |
| Cell dimensions | |
| *a*, *b*, *c* (Å) | 67.79, 100.03, 103.25 |
| α, β, γ (°) | 90.00, 90.00, 90.00 |
| Resolution (Å) | 29.34 – 1.90 (1.97 – 1.90)[a] |
| $R_{merge}$ | 0.21 (1.75) |
| $I / \sigma I$ | 9.7 (1.5) |
| $CC_{1/2}$ | 99.7 (65.6) |
| Completeness (%) | 99.1 (98.3) |
| Multiplicity | 13.8 (14.0) |
| **Refinement** | |
| Resolution (Å) | 29.34 – 1.90 |
| No. reflections | 55,515 (5,407) |
| $R$ / $R_{free}$ | 0.180 / 0.218 |
| No. non-H atoms | |
| Protein | 4719 |
| Ligand | 15 |
| Water | 356 |
| *B*-factors | |
| Protein | 32.72 |
| Ligand | 39.75 |
| Water | 42.02 |
| R.m.s. deviations | |
| Bond lengths (Å) | 0.007 |
| Bond angles (°) | 0.83 |
| Ramachandran plot statistics | |
| No. favored | 598 (98.17 %) |
| No. allowed | 11 (1.83 %) |
| No. outliers | 0 (0 %) |

Data set was collected from a single crystal. [a]Values in parentheses are for highest-resolution shell.

**Table S1.** Data and refinement statistics for crystal structure of SARS-CoV-2 main protease ($M^{pro}$) in complex with peptide from human tRNA methyltransferase TRMT1 (PDB 9DW6).

| TRMT1 Co-Incubation | $k_{obs}$ (hr$^{-1}$) | $k_{obs}$ +/- | $K_D$ (µM) | $K_D$ +/- |
|---|---|---|---|---|
| $M^{pro}$ WT | n.d. | n.d. | 4.76 | 1.08 |
| $M^{pro}$ C145A | 1.97 | 0.07 | 0.86 | 0.03 |
| No Protease | 2.10 | 0.05 | 0.80 | 0.05 |

**Table S2.** Methyltransferase activity ($k_{obs}$) and tRNA binding affinity ($K_D$) parameters for TRMT1 after an 18-hour incubation with $M^{pro}$ WT, C145A, or no protease, corresponding to the fits of plots presented in Figure 2C and D, respectively. TRMT1 tRNA modifying activity was measured by radiolabel-based methyltransferase assays with *S*-[methyl-$^{14}$C]-adenosyl methionine and fit to a first-order exponential to obtain $k_{obs}$ (see also Figure 2C); TRMT1 incubated with $M^{pro}$ WT did not have activity during the 4-hour time course, so $k_{obs}$ is listed as n.d. (not determined) for this condition. TRMT1-tRNA binding affinity was determined by EMSA (see **Figure S2**) and fit to a standard single-site ligand binding equation to obtain $K_D$s (see also Figure 2D). All kinetic and binding experiments were carried out in triplicate and errors are reported above as the standard error of the fits shown Figure 2C and D.

| Substrate | $M^{pro}$ Mutation | [$M^{pro}$] (µM) | $k_{cat}$ (s$^{-1}$) | $k_{cat}$ +/- | $K_M$ (µM) | $K_M$ +/- | $k_{cat}/K_M$ (µM$^{-1}$s$^{-1}$) | $k_{cat}/K_M$ +/- |
|---|---|---|---|---|---|---|---|---|
| nsp4/5 | WT | 0.05 | 1.0492 | 0.04640 | 109 | 7.8 | 0.0097 | 0.00026 |
| nsp4/5 | M49A | 0.05 | 0.4142 | 0.02900 | 75 | 9.6 | 0.0055 | 0.00032 |
| nsp4/5 | N142A | 0.05 | 1.0632 | 0.07180 | 122 | 12.9 | 0.0087 | 0.00033 |
| nsp4/5 | Q189A | 0.05 | 0.5274 | 0.10260 | 86 | 29.2 | 0.0061 | 0.00088 |
| nsp8/9 | WT | 0.05 | 0.0190 | 0.00159 | 40 | 7.5 | 0.0005 | 0.00005 |
| TRMT1 | WT | 0.05 | 0.0052 | 0.00024 | 29 | 3.2 | 0.0002 | 0.00001 |
| TRMT1 | M49A | 0.05 | 0.0064 | 0.00030 | 24 | 2.9 | 0.0003 | 0.00002 |
| TRMT1 | N142A | 0.05 | 0.0081 | 0.00038 | 31 | 3.5 | 0.0003 | 0.00002 |
| TRMT1 | Q189A | 0.05 | 0.0062 | 0.00048 | 38 | 6.7 | 0.0002 | 0.00002 |
| TRMT1(A531S) | WT | 0.05 | 0.0040 | 0.00069 | 22 | 8.3 | 0.0002 | 0.00004 |

**Table S3.** Michaelis–Menten kinetics determined for different fluorogenic peptide substrates cleaved by $M^{pro}$ wild-type (WT) and $M^{pro}$ mutants (M49A, N142, Q189A). Individual kinetic measurements were carried out in triplicate and the +/- columns denote the standard errors on each parameter derived from non-linear least squares regression fits.

**Dataset S1 (separate file).** Peptide cleavage kinetic assay initial rates measured for $M^{pro}$ WT and $M^{pro}$ variants with nsp4/5, nsp8/9, TRMT1, and TRMT1A531S substrates, used to determine the kinetic parameters published in this manuscript.

**Dataset S2 (separate file).** TRMT1 rodent and primates orthologous sequences used for evolutionary analysis and positive selection analysis of rodent and primate TRMT1 genes.
